# Has recombination changed during the recent evolution of the guppy Y chromosome?

**DOI:** 10.1101/2023.05.17.541215

**Authors:** Deborah Charlesworth, Suo Qiu, Jim Gardner, Roberta Bergero, Karen Keegan, Abigail Hastings, Mateusz Konczal

**Affiliations:** Institute of Ecology and Evolution, School of Biological Sciences, University of Edinburgh, Charlotte Auerbach Road, Edinburgh EH9 3LF, U.K.; Scottish Rural Agricultural College, Peter Wilson Building, King’s Buildings, W Mains Rd, Edinburgh EH9 3JG; Evolutionary Biology Group, Faculty of Biology, Adam Mickiewicz University; 60-614 Poznań, Poland

**Keywords:** Partial sex linkage, pseudo-autosomal region, sexually antagonistic polymorphism, sex reversal

## Abstract

Genome sequencing and genetic mapping of molecular markers has demonstrated Y-linkage across most of the guppy (*Poecilia reticulata*) XY chromosome pair. However, they also revealed exchanges with the X, consistent with classical genetic observations of occasional exchanges of factors controlling visible male-specific phenotypic traits. It remains unclear whether this fish species has an extensive sex-determining region without crossing over (whose suppressed recombination could have evolved under selection created by sexually antagonistic effects of male coloration factors), or whether the fully Y-linked region is very small, perhaps within a single gene. Population genomic data support cytogenetic results indicating that it is within the terminal 5 Mb of the 26.5 Mb chromosome 12, just proximal to a highly recombining pseudo-autosomal region, PAR1. Using molecular markers, we studied recombination, focusing on this region of the XY pair. Despite assembly errors in the small terminal PAR1, our mapping identifies very similar genetic PAR boundaries in sires from four populations, suggesting that their crossover patterns have not changed. Our results also confirmed occasional crossovers proximal to the male-determining region, defining a second pseudo-autosomal region, PAR2, recombining much more rarely than PAR1. The crossover positions suggest that the male-determining factor is within a repetitive region near 21 Mb in the female assembly. A sex-reversed XX male had few crossovers in PAR2, suggesting that this region’s low crossover rate depends on the phenotypic, not the genetic, sex. Thus rare sex changes, and/or occasional crossovers in males can explain the failure to detect fully Y-linked variants.

## Introduction

Guppy (*Poecilia reticulata*) populations have been important for studies relating to adaptive evolution and the evolution of sex chromosomes, especially the rarity or absence of recombination between the Y- and X-linked regions. More than 100 years ago, the guppy Y chromosome of this sexually dimorphic fish was shown to carry a male coloration factor (Schmidt 1920), and subsequent studies found several more such factors, many of them identified in domesticated strains, but also one controlling the naturally polymorphic maculatus pattern (reviewed in Winge 1927). Such coloration is associated with higher predation rates, but frequency-dependent advantages in males (rare coloration phenotypes gain matings and survive better than ones at higher frequencies), allow alleles of these factors to be maintained as polymorphisms at intermediate frequencies (Haskins and Haskins 1951; Potter *et al*. 2023). The most comprehensive study of naturally polymorphic male coloration “patterns” (Haskins and Haskins 1951; Haskins *et al*. 1961). confirmed Winge’s conclusions based on genetic results for 312 traits found in Trinidad rivers (contributing to 8 trait combinations named by earlier workers and listed in Table 11 of Haskins, 1961).

Using both father-to-son transmission (44 sires and 749 sons), and trait expression in 537 female progeny treated with testosterone to allow female carriers of the male-specifically expressed coloration factors to be detected, 75% of the patterns were classified as showing Y-linked transmission, while only 10% were autosomal (summarised in Table 12 of Haskins *et al*. 1961). Given that the guppy has 23 chromosomes, and that the sex chromosome, chromosome 12, represents less than 5% of the species’ genes (Künstner *et al*. 2017), the Y chromosome is strongly over-represented. Other patterns showed evidence for partial sex linkage. Winge (1934) also found one recombinant between the factors controlling the red and the black elements of the maculatus pattern among 3,800 progeny studied, and several later genetic studies confirmed such recombination. In one multi-factor linkage map, the variegated tail factor (*Var*) is on one side of the sex-determining locus, and genes for 10 other visible traits are within < 15 cM on the other side, suggesting the presence of two pseudo-autosomal regions (Khoo *et al*. 1999). Lindholm and Breden (2002) reviewed all available studies, including those in domesticated guppies, showing that Y-X recombination occurs, albeit at low rates, and also that females carrying coloration factors do not express many coloration traits, potentially explaining the male-limited coloration phenotypes in natural guppy populations. This early evidence that recombination is not completely suppressed in the guppy is confirmed by the finding that its Y chromosome has not undergone genetic degeneration (Bergero *et al*. 2019; Fraser *et al*. 2020; Almeida *et al*. 2021; Charlesworth *et al*. 2021).

Nevertheless, both the predominance of Y-linkage, and male-specific expression, suggest that the factors controlling male coloration polymorphisms also have sexually antagonistic (SA) effects that reduce fitness in females. If the mutations that created the male coloration factors showed male-specific expression when they first arose, nothing would prevent their spread in populations even if they were autosomal, and thus over-representation on the Y chromosome would not be expected.

Y-linkage of male coloration factors thus suggests preferential spread of mutations that arose in partially sex-linked genes that recombine infrequently with the male-determining locus. This is an example of a “selective sieve” in which the fate of a favorable mutation depends on linkage to a balanced polymorphism, with mutations in genomic locations closely linked to a previously established polymorphism having a lower rate of loss from the population than mutations with the same phenotypic effect that arise in other, less closely linked, locations (Charlesworth and Charlesworth 1975; Turner 1977; Charlesworth and Charlesworth 1978). Male coloration mutations that become polymorphic within guppy populations should therefore cluster within a genome region closely linked to the male-determining locus, satisfying the “sieve” criteria (Jordan and Charlesworth 2012). Male-specific expression (or suppressed expression in females) would then be favoured for such polymorphic factors, because expression of the traits is detrimental in the occasional recombinant females. This scenario can therefore explain both the predominance of Y linkage, and the male-limited expression, of guppy male coloration factors. However, the evolution of expression changes awaits testing.

It also remains unclear whether closer linkage to the guppy male-determining factor has evolved, or whether, as the “sieve” hypothesis proposes, the present recombination rates and patterns evolved before the polymorphic coloration factors became established (Bergero *et al*. 2019; Charlesworth *et al*. 2020b). Although polymorphic SA factors are of special interest, because they can create selection for both sex-limited expression and/or closer linkage between the two loci involved (in this case, the sex-determining locus and the gene with the polymorphic SA alleles (Rice 1984; Rice 1992), and can be investigated by genetic analyses, use of these traits in genetic studies could not resolve this question. Indeed, the predominance of Y-linkage has recently been questioned, based on analyses of individual orange and black spots in a captive population studied derived from a high-predation site of the Quare river in Trinidad (Morris *et al*. 2020). These authors concluded that loci controlling the presence or the absence of individual male traits are not predominantly Y-linked. Excluding traits shared by almost all sires, which are uninformative for genetic study, and tail traits (which are often not Y-linked), one orange spot, OI, showed largely Y-linked inheritance, while three black spots were often not controlled by simple major Mendelian Y-linked factors. However, large grandpaternal effects were found for the composite orange and black spot area phenotypes (whereas grandmaternal effects were mostly negative), suggesting a strong paternal transmission component (especially given that non-variable traits were included). This is consistent with other studies reviewed by MORRIS et al. (2020), including previous inferences that modifiers are involved (Haskins and Haskins 1951; Tripathi *et al*. 2009), and with analyses using guppies collected in Australia, which estimated that about half of the genetic effects represent Y-linkage (Postma *et al*. 2011). Overall, therefore, an unexpectedly large Y-linked effect appears to be convincingly demonstrated. However, different Y-linked, factors that recombine very rarely, at most, cannot be individually identified by classical genetic approaches, so it remains unknown how many individual male coloration factors are carried on the guppy Y chromosome, or what the factors are, or even rough locations on the chromosome (see the Discussion section).

As will be described shortly, there is evidence that recombination may differ between different guppy populations, which would suggest ongoing evolution of crossover patterns and/or rates. Given the uncertainties just mentioned concerning the genetics of coloration factors, however, it would be preferable to estimate rates using molecular markers, whose genetics is much clearer. Combined with physical mapping, using known chromosomal locations of markers, one can test whether guppy populations differ in recombination. If not, the observed Y linkage may not reflect effects of SA selection, and seems more likely to reflect a sieve acting during establishment of the male polymorphisms, along with recombination patterns that had evolved before the establishment of extant populations.

Long before the availability of genome sequencing, evidence for linkage disequilibrium (or LD) was described in natural Trinidadian guppy populations between at least some coloration factors and the male-determining locus, and evidence that guppy populations from Trinidad rivers might indeed differ in recombination. The results were based on the fact that, despite their male-specific expression, genetic coloration factors can be detected in females by testosterone treatment. This consistently revealed higher frequencies of female carriers of several coloration factors in upstream than downstream populations (page 374 of Haskins *et al*. 1961). For the widely distributed Sb coloration polymorphism, whose frequencies are generally somewhat below 20% (Table 11 of Haskins *et al*. 1961), a sample of 33 males from a downstream population in the Aripo river (with high predation) transmitted the Sb factor almost exclusively to male progeny, and not their female progeny, demonstrating Y-linkage. Of 19 Sb males from a low-predation site, however, only 8 transmitted the trait exclusively to sons, while four transmitted it to nearly all their female progeny; thus these males’ X chromosomes carried the factor. Seven other males also carried the factor on both the X and Y (all their male and testosterone treated female progeny expressed the Sb phenotype). The *Sb* factor is therefore almost completely Y-linked only in the downstream population (Table 14 in Haskins *et al*. 1961), but in the upstream male parental sample, the Sb factor was X-linked almost as often as it was Y-linked (in 11 versus 15 of the 19 sires, or 58 versus 79%, respectively). Additionally, only a single potentially recombinant individual was found among 1,024 progeny of downstream males, versus 4 among 387 progeny of upstream males (a 13-fold difference in the proportions, P = 0.0071 in a two tailed Fisher’s exact test).

Similar observations of associations with sex (without genetic mapping results in sibships) were obtained for factors controlling the proportions of females that developed black or orange coloration under testosterone treatment in samples from the Aripo and four other rivers; results from a low-predation Paria river sample were similar to those from low-predation sites in other rivers (Gordon *et al*. 2012, see Figure 1). Estimated frequencies of different polymorphic male coloration factors differ between collection sites in Trinidad (Table 11 of Haskins *et al*. 1961). Frequency data are, however, extremely scarce, in part due to the difficulty of determining the genetics of these factors (see above). Individual allele frequencies cannot therefore be studied. The more than 20% of females with factors for orange and black spots detected by Gordon *et al*. (2012), even in high-predation populations, probably reflect both autosomal and sex-linked factors, both of which must be male-limited in expression, but cannot be distinguished.

**Figure 1.**
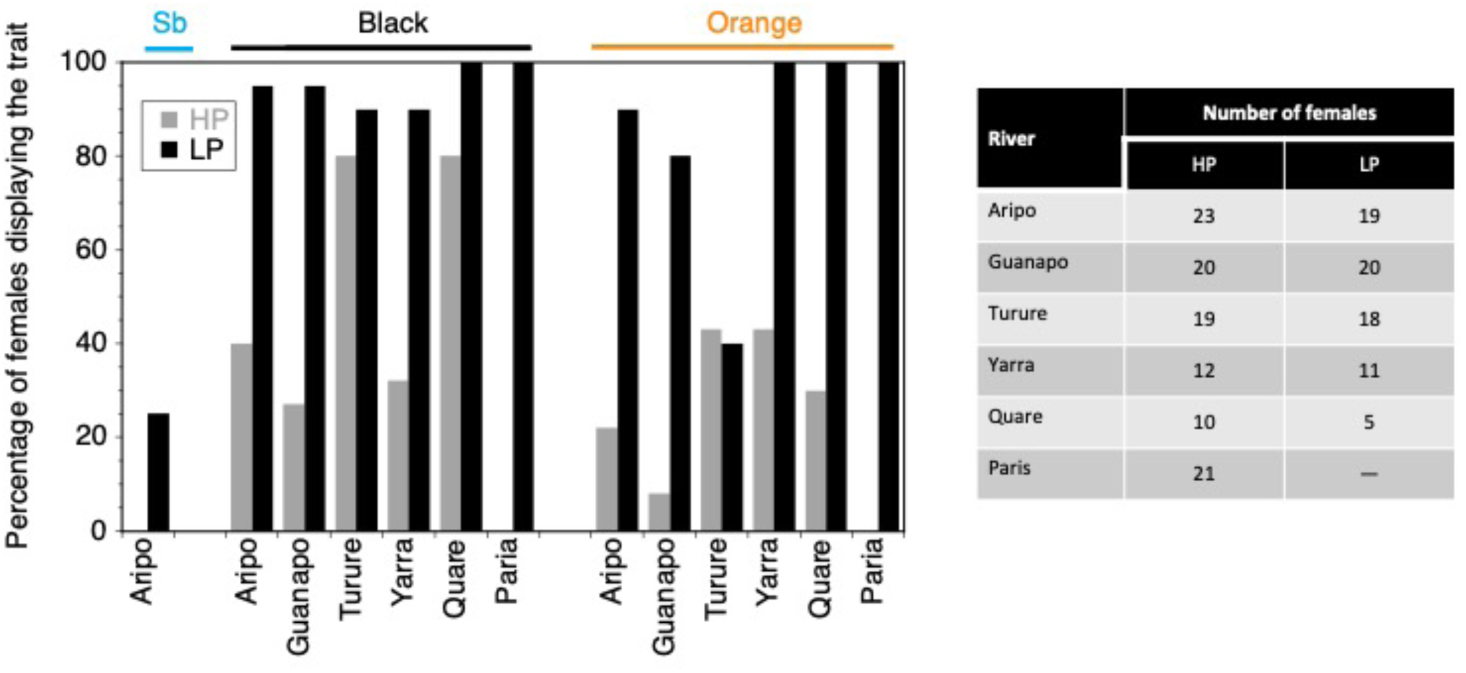
Summary of the results from studies of associations between individuals’ phenotypic sexes and coloration in the experiments of Haskins (1961) and Gordon et al. (2012) outlined in the main text.

Consistent with the genetic results above suggesting that these composite phenotypes are controlled by multiple loci, not all Y-linked (Morris *et al*. 2020), some individual traits are fixed in the natural population samples, as all testosterone-treated females expressed the male coloration phenotype (see Figure 1). The results from females from low-predation populations nevertheless consistently indicate that coloration factors are often autosomal or X-linked in such populations (Gordon *et al*. 2012). Fixation could reflect bottlenecks at their foundation, allowing loss of low frequency coloration alleles by genetic drift, and population genomic results support the occurrence of recent bottlenecks affecting up-river populations in Trinidad (Qiu *et al*. 2022). Alternatively, net selection against males showing such traits may be weaker in upstream environments. The striking difference from the results for the Sb factor probably reflects the simple Y-linked inheritance for this non-composite phenotypic trait.

Overall, the available results strongly suggest that, in low-predation populations, some coloration factors may be commonest in upstream, low predation sites, but that coloration genes show Y-linkage more often in high-predation populations. There could also be genetic variation affecting the expression of coloration traits in testosterone treated females. Autosomal factors suppressing expression in females carrying such factors might, for example, be rarer in upstream populations than in high-predation ones. It is thus uncertain whether recombination rates between coloration factors and the sex-determining locus in male meiosis differ between high- and low-predation guppy populations, or whether the associations of coloration factors with maleness differ for other reasons.

Here we directly test for differences in the genetic maps from males from upstream and downstream populations from the Aripo and Quare rivers, using molecular markers (microsatellites and single nucleotide polymorphisms). The guppy sex chromosome pair shows strong crossover localisation in male meiosis, with almost all crossovers being found in the terminal 5% of this telocentric chromosome (Bergero *et al*. 2019; Charlesworth *et al*. 2020a), which is a highly recombining pseudo-autosomal region (labelled PAR1 in Figure 2). Consistent with the results for the *Sb* factor outlined above, few recombinants with the sex-determining locus are detected proximal to the PAR1 boundary, at about 25 Mb in the female assembly. There is thus a second PAR (PAR2), which is very different from PAR1, as crossovers in PAR2 are rare. However, only the unknown male-determining factor itself (labelled M in Figure 2) is definitively known to be completely Y-linked, but neither its location, nor the extent of any surrounding fully Y-linked region, are yet known (see below).

**Figure 2.**
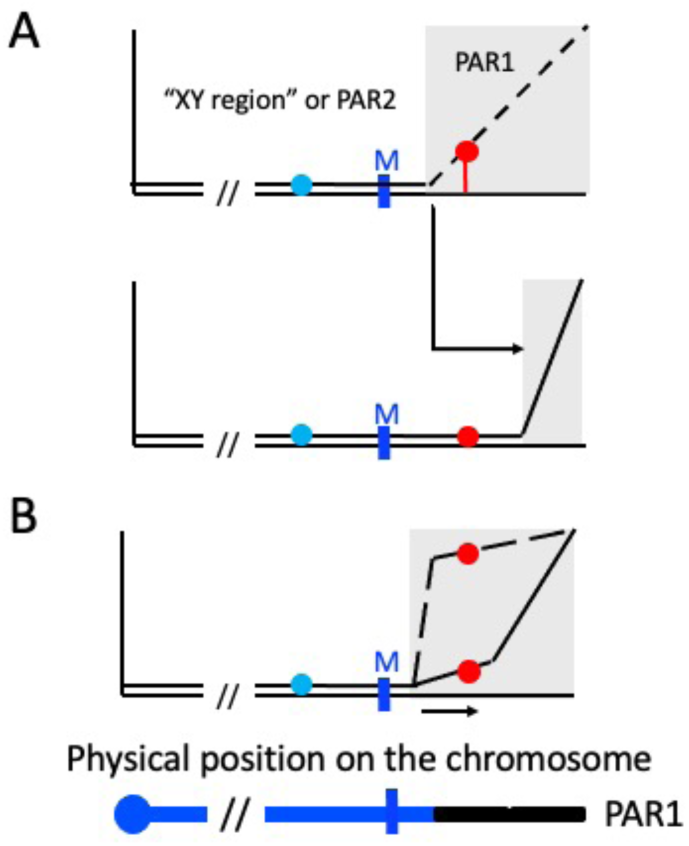
Diagrams of situations that could create differences in recombination rates in male meiosis between coloration factors and a male-determining locus. The organisation of the guppy sex chromosome is diagrammed at the bottom, with the centromere at the left, and a male-determining factor in the terminal third of the chromosome (indicated by a dark blue vertical bar labelled M). Recombination rates with the M factor are indicated by black lines. The M factor is proximal to a pseudo-autosomal region that recombines at a high rate with the male-determining region, termed “PAR1” and indicated by grey boxes. The region to the left of PAR1 is termed the “XY region” since it segregates as almost completely sex-linked (the lines indicating the recombination frequency with the M factor are close to the x axis). However, crossing over occasionally occurs, so this region could also be termed “PAR2”. Male coloration factors closely linked to the M factor could be located in the XY region or in PAR1 near the boundary with the XY region (the diagrams indicate hypothetical factors by blue and red circles). **A**. Two coloration factors both initially linked to the maleness factor, M. Movement of the boundary to the right, by an inversion or other crossover modifier, causes the factor in the more highly recombining PAR1 to be within an XY region gene, and PAR1 becomes physically smaller, and the recombination rate distal to the boundary is high, as in extant guppy populations. **B**. Change of the crossover distribution within PAR1. The initial pattern, shown by a dashed line, changes to move crossovers away from the boundary with the XY region (the unbroken line), creating closer linkage between the red and M factors.

A lower recombination rate in the downstream populations (with strong associations between male coloration and the Y-linked maleness factor) could thus be due to different locations of the PAR1 boundary, such that a partially sex-linked region in upstream males is fully sex-linked in downstream ones (A in Figure 2), or to a difference in the recombination rates within PAR1 (Figure 2B), or both. We used locations in the guppy female and male assemblies (Künstner *et al*. 2017; Fraser *et al*. 2020) to relate map locations in male and female meiosis to the physical locations of genetic markers and test between these possibilities using the male maps. Previous genetic maps from F2 individuals (Whiting *et al*. 2021) could not separate male from female meiosis, and the roughly constant recombination rate in female meiosis across the entire chromosome obscured the relationships between genetic and physical positions. This previous mapping did, however, support elevated recombination rates in male meiosis near chromosome termini (including the sex chromosome’s PAR1).

## Methods

### Families used for genetic mapping

Supplementary Table S1 lists the sources of the parents, all of which were captured in the natural populations indicated in the table, and the numbers of parents and progeny used for studying recombination. Detailed maps were estimated for two Aripo river families, and two families from the Quare river, in each case from one high- and one low-predation site (indicated by H or HP in the family name for the former, and by LP for the latter, see Table S1). Crosses were made between the parents after these were transported to the University of Exeter, Falmouth, except for the family from the Aripo high-predation site, whose parents were from a captive population that had been maintained for several years since being collected (Bergero *et al*. 2019; Charlesworth *et al*. 2020a). All the families except this one consisted of several full sibships with different parental individuals. In sibships with multiple possible parents, parentage of progeny was inferred using genotypes at multiple microsatellite markers and confirmed using SNPs in the high-throughput genotyping data (see below).

### Genetic markers

SNPs for genetic mapping were genotyped in high-throughput experiments performed by LGCG, https://www.lgcgroup.com/ (Charlesworth *et al*. 2020a). The targets were selected in coding sequences annotated in the guppy female genome assembly (Künstner *et al*. 2017), using NCBI Poecilia reticulata 1.0: https://www.ncbi.nlm.nih.gov/assembly/GCF000633615.1/. For our first genetic mapping experiment, Experiment 2, a set of about 500 target sequences were selected from the sex chromosome pair, and smaller numbers from each of the 23 guppy autosomes. Experiment 3 used a larger number of SNPs from the sex chromosome pair, and Experiment 4 mapped the same set of SNPs from the sex chromosome pair, plus large numbers from the autosomal chromosome 16. The male genome assembly is not annotated, and the male assembly positions were found by BLAST searches, which identified most, but not all, sequences targeted (See Supplementary Tables 3C and D). For sex-linked targets that were not assembled on chromosome 12 in the male assembly, microsatellite markers were designed for mapping in Experiment 2 (Supplementary Table 2B).

Data pre-processing by LGCG included demultiplexing of all libraries for each sequencing lane using the Illumina bcl2fastqv2.20 software [Illumina. bcl2fastq2 Conversion Software. URL: https://support.illumina.com/sequencing/sequencing_software/bcl2fastq-conversion-software.html] (folder ‘RAW’ subfolders), allowing at most 2 mismatches or Ns in the barcode read when the barcode distances between all libraries on the lane allowed for it. The sequencing adapter remnants were then clipped from all reads (folder “AdapterClipped”) and reads with final lengths < 20 bases were discarded before merging forward and reverse reads using BBMerge v34.48 citebbtools (folder “Combined”); the consensus sequence of combinable fragments are named “*joined-SR*”, and sequences with read pairs that could not be combined were stored in files named “*R1*” and “*R2*”. The remaining (combined) reads were quality cropped, setting the quality of the first 40 bps (corresponding to the synthetic oligonucleotide probe) to 0 to avoid biased genotyping in the downstream analysis, and the resulting sequences are in the “QualityTrimmed” folder. FastQC reports were created for all FASTQ files (Andrews 2010). Finally, read counts for all samples were generated. This involved initial alignment of subsampled quality trimmed reads against the reference sequence using BWA-MEM v0.7.12, http://bio-bwa.sourceforge.net/ (Li 2013), followed by variant detection and genotyping of all samples using Freebayes v1.2.0, http://github.com/ekg/freebayes (Garrison and Marth 2012), with ploidy set to 2, since the guppy Y chromosome is not genetically degenerated. Genotypes were filtered to a minimum coverage of 8 reads.

Microsatellite sequences were found in the guppy female and male genome assemblies (Künstner *et al*. 2017; Fraser *et al*. 2020) using the PERF software v0.4.5 (Avvaru *et al*. 2018). Excel files listing microsatellites for all guppy chromosomes, based on both the male and female assemblies, including the PCR primer sequences for amplifying them, and files with the marker locations and the genetic mapping results have been deposited in Figshare.

### Genetic mapping

The high-throughput genotyping experiments produced counts of each allele inferred to be present in each individual from the chromosomes targeted. Duplicate DNA samples of some individuals were included, to assess the reliability of the results. These files, with the marker locations, and the genetic mapping results are also available in Figshare.

The marker results were analysed using the LepMap3 software package, which is designed for mapping data from families consisting of half-sibships with shared parents (Rastas 2017). The genotypes were analysed as autosomal markers, because the guppy Y chromosome is not genetically degenerated, and appears to carry all genes present on the X (Bergero *et al*. 2019; Fraser *et al*. 2020; Almeida *et al*. 2021; Charlesworth *et al*. 2021). Before estimating genetic maps, the total allele counts for all individuals in a family to be analysed data were filtered to exclude sequences with low coverage (for heterozygous individuals, the counts for both alleles were used); these counts are in the files deposited in Figshare. The duplicate with the higher total count across all sequences genotyped was then used for the mapping analysis. Further filtering within each family was done by LepMap3, to combine individuals with duplicate sequences, and using the parameters missingLimit=0.5 to filter out markers with different proportions of missing genotype data, and dataTolerance=0.0001 for the Aripo river data for the P-value threshold for excluding SNPs with non-Mendelian segregation ratios. For the Quare river data (with much larger numbers of markers, see below), a dataTolerance value of 0.04 was used, and an interference value of 0.03 was applied for the QLP1B1 data, and 0.02 for the QHPG5 family. Supplementary Table S3 shows the informative markers called for each guppy chromosome assembly in each family.

Markers from all 23 guppy chromosomes were genotyped in the two Aripo river families. LGs with < 5 markers were removed using sizeLimit=5 for the SeparateChromosomes module of LepMap3. Initial analysis with several values of the lodLimit parameter indicated that a value of 7 gave the closest number of linkage groups (LGs) to the guppy chromosome number for the LAH family data (18 chromosomes formed single LGs, while 5 were divided into 2 separate LGs; a total of 1,104 markers were mapped to these 28 LGs. These families, and two Quare river families, again from a low predation up-river and a high-predation down-river site, were also mapped with dense markers from the sex chromosome, in order to map the terminal PAR1 part of the chromosome in the sires. Chromosome 16 markers were also genotyped in the up-river Quare family (see the Results section).

### Identifying the centromere ends of the guppy chromosomes

In male meiosis, many markers informative for mapping co-segregated in the families, and these identify the end of the assembly that corresponds to the centromere, and were used to identify the zero centimorgan position in the genetic maps. In the dams, many markers were not informative for mapping, and the most centromere-proximal markers in the genetic maps for female meiosis are not always physically near the centromeres (see Supplementary Table S3). The estimated female map lengths for several chromosomes mapped in the two Aripo families in Experiment 2 (with fewer markers mapped than in Quare river families in Experiments 3 and 4) are therefore shorter than their true lengths.

To further check the centromere ends of the chromosomes, we calculated GC content, which is expected to increase at the telomere ends due to GC-biased gene conversion. An analysis using the female assembly based on short sequencing reads detected the predicted effect, consistent with high recombination rates in guppy male meiosis in these regions (Charlesworth *et al*. 2020b). We repeated the previous analysis using the male assembly, which is preferable, as it is based on long-read sequencing (Fraser *et al*. 2020). Also, to avoid possible enrichment with GC-rich transposable elements near the chromosome termini, we masked repetitive sequences. After using RepeatModeler2 (Flynn *et al*. 2020) to generate an initial comprehensive set of candidate TE families in the male assembly. RepeatMasker (Smit et al., http://repeatmasker.org) was then used to make a library of consensus sequences from the RepeatModeler2 analysis plus Teleostei repeats from Repbase. A filtered list of candidates was created, and manually annotated, following the procedures and criteria described by Goubert *et al*. (2022). After obtaining candidate sequences, RepeatMasker was run again to find and annotate repeat copies throughout the male reference genome. Files containing the consensus repeat library and localization of repeats in reference genome were deposited in the Figshare repository. Finally, GC content was estimated in 20kb non-overlapping windows using a custom script that excluded the annotated repeats and sites in the reference assembly other than A, T, G or C.

## Results

### Crossover localisation in guppy chromosomes: genetic maps of the autosomes

Female and male genetic maps from families were estimated using high-throughput genotyping of parents collected from high- and low-predation sites from the Aripo and Quare rivers and their progeny (details of the sources of the samples, and the experiment numbers and progeny sizes are in Supplementary Table 1). Female maps from two Aripo river families confirm the previous conclusion using sparse microsatellite markers (Bergero *et al*. 2019) that crossovers occur across the whole of each guppy chromosome. Several chromosomes could not be reliably mapped in the low-predation sibships from the Aripo river, due to insufficient SNPs. Whenever maps could be estimated, however, they support the conclusion that crossovers occur evenly across each chromosome, including the X (see Figure 3 in the next section). The results also confirm that the recombination pattern is very different in males (Charlesworth *et al*. 2020b). In both families, our maps detect an extensive region at one end of each chromosome (usually <u>at least</u> 80% of the assembly) consistently lacks recombinants in male meiosis. Chromosome 16 had previously been a possible exception to this, so we attempted to map it using SNPs genotyped in the QLP1B1 family in experiment 4. Supplementary Figure S3 shows that, in male meiosis, crossovers are probably localised at the end of this chromosome.

**Figure 3.**
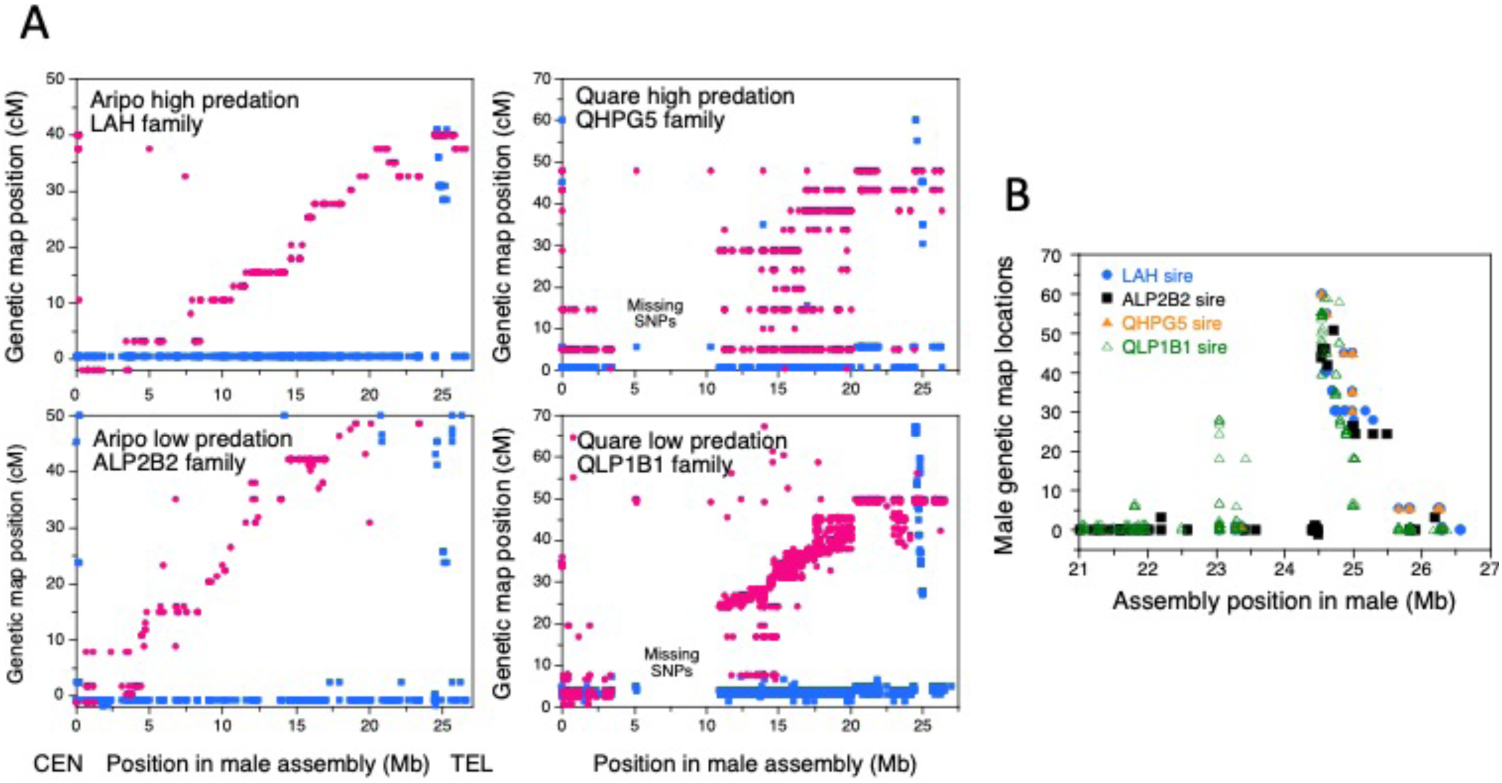
Genetic map results from parents of the four families indicated. A. Male and female maps for the whole of the sex chromosome pair. B. Maps from the male parents of the families indicated in the key, for the terminal part of the guppy sex chromosome, starting from 21 Mb in the male assembly. Markers proximal to about 24.5 are not part of the highly recombining pseudo-autosomal region, PAR1, in any of these families, which all have very similar PAR1 boundary positions.

In both female and male meiosis, there is generally one crossover per bivalent, though some chromosomes lack markers informative in male meiosis in the terminal region, and have much smaller map lengths. Total genetic map lengths of about 50 cM in male meiosis are consistent with the cytologically estimated one crossover per bivalent (Lisachov *et al*. 2015).

Markers informative in the dams were not always available across the complete physical assembly of all 23 guppy chromosomes, so the ends are not always mapped (Supplementary Figure S1). Some chromosome maps are therefore smaller than 50 cM, and the locations in female meiosis may be lower than the correct values. We therefore did not compare the map lengths in the two sexes. The regions in which crossovers are infrequent in the male maps identify the genetical centromere ends. These are at the start of the assembly for all except 6 chromosomes (Supplementary Table S2), consistent with inferences from elevated GC content in introns and third codon positions, based on the reasoning that regions with very high recombination rates in males should have elevated GC content, due to GC-biased gene conversion (Charlesworth *et al*. 2020b). As it is possible that this pattern might instead reflect preferential presence of GC-rich transposable element(s) near the chromosome termini (Schield *et al*. 2022), we repeated the analyses of GC content after masking repetitive sequences ascertained using RepeatModeler, as described in the Methods section (Flynn *et al*. 2020). This found the same centromeric ends as those previously inferred (Supplementary Figure S2).

When SNPs near the centromere ends could be mapped in female meiosis, the results for multiple autosomes appear to have large regions with few recombinants, if any, in female, in both Aripo river families (including chromosomes 6, 7 10, 11, 13, 16, 16, 20 and 21). If population genomic results find low synonymous site diversity in these regions, this would suggest that the guppy may have extensive pericentromeric regions with low crossover rates like those in many other eukaryotes, in which centromere effects on recombination are common reviewed in Miller and Hawley (2017). There is no sign of any low recombination region on chromosome 2, despite the fact that it arose by a centric fusion that created a single chromosome from homologues of the platyfish chromosomes Xm2 and Xm24 and is physically large (Künstner *et al*. 2017), see Supplementary Table S2, and has a longer genetic map than other chromosomes.

### Crossovers on the guppy sex chromosomes and the PAR1 boundary in our families

Figure 3A shows female and male maps of the guppy sex chromosome (12) in sires from two Aripo and two Quare river families, again using high-throughput genotyping, but targetting 500 guppy sex chromosome sequences, in an effort to map as much of this chromosome’s assembly length as possible. Map locations close to zero should therefore be close to the correct values (unlike the situation for the autosomes). The female meiotic maps can therefore be compared between the high- and low-predation families, and are clearly very similar.

In the female assembly, a region between 10 and 20 Mb is inverted, probably reflecting an assembly error that was corrected in the male assembly (Fraser *et al*. 2020). The new maps support our previous conclusion for female meiosis, that the guppy X chromosome arrangement is always the same as that in the male assembly (though the family from the Quare low-predation population includes only 23 progeny, and the female map is poorly estimated). The region near 10 Mb in the female assembly is repetitive, and these genes are not in the assembly of the platyfish homologue, Xm8. SNPs were available in 9 of these genes (3 assigned to chromosome 17 in the male guppy assembly, one assigned to chromosome 15, and 5 to chromosome 7) and all segregated autosomally, some in several families (Supplementary Table S3B, C and D). The chromosome 7 genes include the Olr1496-like gene with a function in reproduction that shows higher coverage in guppy males than females, and a high F_ST_ value between the sexes in a guppy natural population sample (Lin *et al*. 2022).

In male meiosis, all crossovers detected are distal to 24.53 Mb in the male assembly, and all more centromere-proximal markers behaved as completely sex linked in all the families studied (Figure 3B); we term this the “XY region” (detailed segregation results for the four families in Figure 3 are shown in Supplementary Tables S3A to D, and Supplementary Table S4 summarises results from all families with results for microsatellite and/or SNP markers distal to 20 Mb in either the male or female assemblies).

### Y-X recombinants

As outlined in the Introduction, the XY region appears not to be completely sex linked, but constitutes a second PAR, or PAR2 that occasionally recombines, but much more rarely than the PAR1 evident in Figure 3. Two recombinant individuals were detected previously (Charlesworth *et al*. 2020a), and we have now found a further family with a recombinant male, for a total of three crossovers in the XY (or PAR2) region among 669 progeny.

By analyses of many more SNP markers than in the previous study, and combining data from the three recombinant individuals, we localised the male-determining factor to a small interval in the female assembly, which is annotated and suggests which genes are in the region identified (the genes are not annotated in the male assembly, and they appear in several different places). Figure 4 summarises the results, which suggest, surprisingly, that all three crossover events occurred near 21 Mb in the female assembly. The recombinant male in the QHPG5 family appears to have two further crossovers in more distal locations, suggesting that the region is inverted in this sire (Wall *et al*. 2022). This possible rearrangement does not affect the conclusions below about candidate male-determining genes.

**Figure 4.**
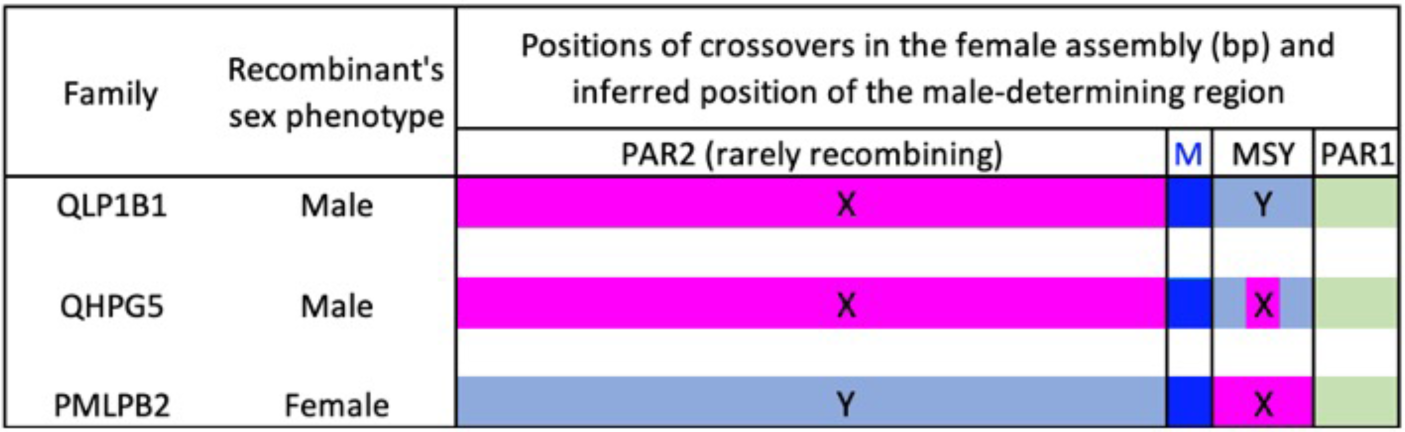
Genetic map results from parents of the four families indicated. A. Male and female maps for the whole of the sex chromosome pair. B. Maps from the male parents of the families indicated in the key, for the terminal part of the guppy sex chromosome, starting from 21 Mb in the male assembly. Markers proximal to about 24.5 are not part of the highly recombining pseudo-autosomal region, PAR1, in any of these families, which all have very similar PAR1 boundary positions.

The new recombinant individual, M7 in sibship 2 of the QLP1B1 family (Supplementary Table S3D), inherited his sire’s paternal X-linked alleles (those found in all 14 female and none of the 6 male full sibs) for 1,660 SNPs informative in male meiosis, from the centromere-proximal site at 130,566 bp until 20,698,741 bp in the female assembly (omitting those that map to chromosome 7 in the male assembly, and 36 SNPs with evidence of duplications or other unexpected segregation patterns); M7’s genotypes change to the sire’s Y-linked alleles at 21,049,564 (or, most likely, from 20,930,721) until the PAR1 boundary (Supplementary Table 3D). The male-determining factor must therefore be distal to 20,698,741 (Figure 4). The recombinant female previously found in the PMLPB2 family (see Supplementary Table 3E) has an almost precisely opposite pattern, inheriting its sire’s X-linked alleles until the marker at 11,748,712 bp in the female assembly (which is close to 20 Mb in the male assembly after correcting the assembly error), and his Y-linked alleles for three markers in the region that are more distal, and that behave as fully sex-linked in all other progeny. As the first such marker is at 21,363,193 in the female assembly, the male-determining factor must be proximal to this, and therefore within the 664,452 bp interval distal to 20,698,741 bp. This interval includes only 24 genes (Supplementary Table S5).

Twelve of these genes are not in the male chromosome 12 assembly, and those that are present are in several different regions. Of the markers mapped in QLP1B1 dams, the 6 in the ptpn13 gene map near 25 cM, while 19 markers (in 9 other genes) map near 50 cM (Supplementary Figure S4); although only two of these are assembled on chr12, they co-segregate with the male-determining locus in male meiosis, and are therefore not autosomal. Four genes are near 5 Mb in the male assembly, close to Contig IV in the male assembly, a sex-determining candidate region which includes repetitive sequences that are duplicated in the more distal contig XII (FRASER *et al*. (Fraser *et al*. 2020). Markers in three contig IV genes were mapped in our families, as well as a microsatellite marker, chr12cIV_AC618 (the only one that yielded reliable genotypes, out of 11 tested in this region, see Supplementary Figure S5A and B). This microsatellite co-segregated with sex in the sires of all three other families (LAH, ALP2B2 and QHPG5); in female meiosis, it could be mapped in LAH and ALP2B2 dams, but the map positions differed from those of other markers near 5 Mb on chr12, and are near much more distal markers. In the QLP1B1 family, SNPs in genes near 5 Mb on chr12 were not genotyped due to lack of SNPs, though, as in other maps, markers in the Contig IV gene cyclin 1 map near 50 cM.

### The sex chromosome terminal region

Figure 3B shows detailed genetic map results for just the terminal part of the sex chromosome, based on high-throughput SNP genotyping, for the two Aripo river and two Quare river families. Even using the male assembly, which used long-read sequences, including PacBio and Hi-C analyses (Fraser *et al*. 2020), the PAR1 genetic map positions are in the opposite order from the physical locations of the markers in both the male and female assemblies (Supplementary Table S5). Several SNP markers in sequences near the end of the male assembly did not recombine with the male-determining locus in any of the four families in Supplementary Tables S3A to D. Both these findings suggest that the entire region is inverted in both guppy assemblies, though the comparison with the female assembly in Supplementary Figure 5 suggests that these may belong in an XY region proximal to the PAR1 boundary.

In addition, markers in several scaffolds that are unplaced in the female or male assembly co-segregate with sex-linked markers (Fraser *et al*. 2020), and can be assigned terminal positions based on the LAH and/or ALP2B2 families (Supplementary Tables 3A and B and Supplementary Figure S5). This includes scaffolds NW_007615023.1 and NW_007615023.1, which are unplaced in the female assembly, but include genes with homologous sequences near the end of the platyfish Xm8 assembly most distant from the centromeres, and map genetically within PAR1. However, markers in unplaced contigs were difficult to map, owing to duplications and/or low coverage or repetitive sequences (reminiscent of Morgan et al.’s 2019 findings in mouse sub-telomeric regions). One of these markers, 023_AC17, was duplicated in some individuals in the ALP2B2 family (but not in the LAH family).

Given the possible assembly problems, the best information for comparing the precise PAR1 boundary locations in different families comes from the most distal markers where no recombinants were seen in meiosis of individual males in experiments 3 and 4, using a higher density of chromosome 12 markers than experiment 2. Wherever information is available, the PAR1 boundary is in a very similar location, near 24.5 Mb in the male assembly (Figure 3B and detailed genotype results in Supplementary Tables S3A to D). All sires yielded very similar maps in the boundary region, with PAR1 markers in the same order when this could be ascertained by mapping. Supplementary Table S6 summarises results from all the families mapped, and identifies the genes closest to the boundary in the female assembly. In the results from experiments 3 and 4, the map distances between markers within PAR1 and ones that co-segregate with the male-determining locus span about 53 cM in male meiosis, and recombination rates within PAR1 appear roughly proportional to physical distances between the markers (Figure 3B), suggesting that there is no recombination hotspot near the boundary between the “XY region” and PAR1.

### Recombination in an XX male

One family from a Guanapo low-predation site (GLPGrp1 in Supplementary Table S1) produced a large number of progeny with a highly female-biased sex ratio. Genotypes of the parents and progeny for four informative microsatellite markers indicated that 15 progeny derived from one sire (8 males and 7 females), sire 1. The 63 other progeny were all female, and were sired by an individual carrying alleles at all four marker loci that were absent from sire 1, thus uniquely identifying this sire (sire 2). This large all-female sibship indicates that sire 2 was an XX male. His alleles segregated in 1:1 ratios in the progeny of two different dams (36 from one dam, and 27 from the other). Winge (1930) also observed sex changes. The appearance of an XX male allowed us to ask whether crossing over is sex-limited and depends on the sex phenotype of the individual.

We therefore asked whether crossovers occur across the whole chromosome (as described above for other XX individuals, which are usually females), or whether such a phenotypic male displays crossover localisation like genetic (XY) males, with no, or rare, crossovers proximal to 20 Mb. One crossover event was detected in the XX male (sire 2), between markers rgrfAC and AG179 (respectively at 3.78 and 17.36 Mb in the male assembly). As described above (Figure 3) crossovers are not expected in males (Supplementary Table S3). If this XX male had a crossover pattern like that of XX female parents, the map distance between these markers in female meiosis predicts that multiple recombinant progeny should be found. In the ALP2B2 family dams, around 35 cM separates the map positions of the *rgrfAC* and AG179 markers, and in the LAH family results *rgrfAC* maps at 4.5 cM in female meiosis, and AG179 at 26 cM. Thus, assuming that the female map is similar in the female parent of the Guanapo family, we might expect more than 10 recombinants among the 63 genotyped progeny. The observed much lower number suggests that this phenotypic male’s crossover pattern is much more similar to that of XY males. Although the one crossover that we observed in this family is unexpected, it could simply reflect the occasional events that occur in male meiosis within the XY region (see above).

## Discussion

### Crossover patterns in guppy male and female meiosis

Our genetic mapping results support the previous evidence that, specifically in male meiosis, crossovers are highly localized to the ends of guppy chromosomes. As the centromeres are located close to one end of each chromosome (they are telocentric or almost so), crossovers are restricted to at most 20% of the opposite ends of all 23 chromosomes. Similar findings have been reported in some other fish where genome sequencing has allowed genetic maps to be compared with physical maps of the chromosomes, including the Atlantic halibut (Edvardsen *et al*. 2022), though other fish species, such as Atlantic herring (Pettersson *et al*. 2019), show much less pronounced patterns of crossover localization and smaller sexual dimorphism in crossover patterns and rates. Our results using molecular markers also support previous inferences of occasional recombination in XY males, and show that it occurs proximal to PAR1.

Our genetic map results do not definitively answer two interesting questions. The first is whether an extensive fully Y-linked region exists. The XX male individual we report appears to recombines like the XY males. Assuming that XY females would similarly have female recombination patterns, as they do in some frog species that undergo such sex changes (Rodrigues *et al*. 2018), recombination in guppy populations could be explained by occasional sex changes, as Winge (1930) proposed. The presence of rare XY females in guppies (Winge and Ditlevsen 1947), can potentially explain the absence of completely Y-linked markers in population genomic studies, as outlined in the Introduction section.

A second question is whether the crossover localization is more extreme in male guppies for the sex chromosome pair than for the autosomes. Although crossovers are often reliably detected as far as 20% of the physical distance from an autosome end in one or other of the Aripo river families in which markers were genotyped from all the autosomes, these results often differed between the two families (Supplementary Figure S1). For example, chromosome 5 appears to be well mapped in both families, and crossovers were detected for multiple terminal markers in the high-predation family (on the left of these figures), but not in the low-predation family, even for the most terminal markers. As a larger number of progeny were genotyped in the latter, this is probably a real difference between the families, which could be reflect effects of the environmental conditions under which the progeny was raised. If so, it may be difficult to obtain recombination estimates that can be applied generally in this species. For chromosomes 11 and 18, and perhaps 23, terminal crossovers were not detected in the sires of either family.

It is thus difficult to compare crossover localization on chromosome 12, the sex chromosome pair, with that for the autosomes (for which our data also differs, as many fewer markers were genotyped than for chromosome 12). However (unlike the variability in male crossover localization on the autosomes), the boundary shown on the left-hand side of the highly recombining PAR1 in Figure 3 is remarkably clear, and its location is remarkably constant, in all four families with dense enough markers to define it (see also Supplementary Table S3). This suggests that the PAR1-XY region boundary has not changed between the up- and down-stream males from either river. In the pair of families from the Aripo river, we genotyped much larger numbers of progeny from the upstream male, so the failure to detect recombination in the boundary region is not due to a small family size. Our family sizes would not have allowed us to detect small recombination rate differences, such as the difference in the small crossover rates between the *Sb* coloration factor and the male-determining locus described above, but they confirm that the large region that recombines rarely occupies very similar parts of chromosome 12 in upstream and downstream males.

The results in Figure 3 are based on the guppy male assembly, and suggest that PAR1 occupies at most slightly more than 1 Mb of this chromosome. Using positions in the female assembly for the same families, Supplementary Figure S7 and Table S3 suggest a size of about 1 Mb. However, the entire PAR1 appears to be inverted in the male assembly, compared with the genetic map (Figure 3). The terminal markers that co-segregate with the sex-determining locus in our families could therefore either belong to the PAR2, which shows almost complete Y-linkage, or they could be variants in sequences within PAR1, but physically near the PAR1-PAR2 boundary, and therefore closely linked with XY region markers. Thus the PAR1 size could be over-estimated. A size of 1 Mb would represent just under 4% of the X chromosome assembly size (Künstner *et al*. 2017), and the Y size in the current male assembly is very similar to that of the X (Fraser *et al*. 2020). Given the uncertainties about the regions on the guppy autosomes that rarely cross over in male meiosis, it remains unclear whether chromosome 12 has evolved stronger crossover localization. However (unlike the variability in male crossover localization on the autosomes), the boundary at the left-hand side of PAR1 is remarkably clear, and its location is almost identical in all four families with dense enough markers to define it, as summarised in Supplementary Table S6.

Our results support the cytological evidence that the terminal PAR1 usually has a single crossover (Lisachov *et al*. 2015), and confirm the division of the guppy XY pair into two regions, PAR2, with infrequent recombination in male meiosis, and a small terminal PAR1, with very high crossover rates between markers, as even a single crossover implies a high recombination rate, as in other small PARs, such as the human PAR1 (Rouyer *et al*. 1986) and the mouse PAR (Marais and Galtier 2003).

The very high crossover rates make reliable genetic mapping possible within PAR1, and the relationship between the genetic and physical map positions appears to be linear (Figure 3B), suggesting a uniform crossover probability across the region, like that diagrammed in Figure 2A. However, there could be a hotspot next to the boundary with the XY region if the markers assembled most terminally do not belong in PAR1, as Supplementary Figure S7 suggests is possible. This can be resolved only when the physical arrangement can be reliably determined, which may be difficult if the terminal region includes repetitive sequences and/or rearrangements, as in some other species (Morgan *et al*. 2019). A linear relationship predicts that characteristics of sequences within the region that are affected by recombination rates, including the GC content at neutral sites, and the mutation rate, should increase sharply at the PAR1-XY boundary, as observed in the mouse (Marais and Galtier 2003). However, these properties can be described only for windows in the genome sequence, inevitably including sequences under different selective constraints, so that the changes may be obscured. In the guppy, the change is moderately sharp, but does not identify the boundary precisely (Charlesworth *et al*. 2020b).

### XY recombination differences between guppy natural populations

The high PAR1 crossover rates also make it possible reliably to detect the boundary with the adjacent chromosome 12 region, which must be either fully Y-linked, or a rarely recombining PAR2. The conclusion that the boundary is very similar in downstream males and males from upstream sites (where predation pressure is less severe, and coloration is less disadvantageous), appears to conflict with the evidence outlined in the Introduction, suggesting that male coloration traits may show closer linkage to the male-determining locus in downstream than upstream populations. However, apart from the results for the *Sb* factor, which was shown to be Y-linked and to recombine rarely with the male-determining locus (Haskins *et al*. 1961), unambiguous evidence for recombination differences is lacking. Both experiments using testosterone-treated females (Haskins *et al*. 1961); (Gordon *et al*. 2012) observed results consistent with coloration being inherited in a non-Y-linked manner most often in females from upstream sites (Figure 1). However, explanations other than higher recombination are not excluded. As some traits are probably controlled by more than a single factor (see the Introduction), higher frequencies of non-Y-linked factors in upstream populations could also account for the observations. If an autosomal, pseudo-autosomal or X-linked factor became fixed in a population, all or most testosterone-treated females would display the trait, depending on the penetrance. The same applies to a composite trait, such as the proportion of females with a given type of coloration (as studied in these experiments), if one or more component traits is controlled by a fixed allele. This can explain the observations (see Figure 1) that, in several up-river sites sampled by Gordon et al. (2012), almost all treated females express the composite traits of black and orange coloration. Therefore, the fact that smaller proportions of treated females from down-river sites showed such coloration need not indicate looser linkage between Y-linked coloration factors and the male-determining locus. It will be difficult to distinguish between different possibilities, given the difficulty of defining male coloration traits with clear segregation and without major effects of environmental conditions and/or other factors in the genetic background. Our genetic mapping results help to distinguish them, by relying on DNA sequence variants that are free from problems of sex differences in expression or effects of the genetic background to provide information about which parts of the chromosome, if any, differ in crossover rates between upstream and downstream populations.

### Associations between coloration traits and the Y-linked region

Our data on recombination across the guppy sex chromosome pair may suggest the location of the male-determining factor within the XY region (Figure 4, Table 1). Based on the female assembly, the observed X-Y crossovers suggest that it is near 21 Mb, not close to the PAR1 boundary between 25.3 and 25.5 Mb (see above). In the family with the highest marker density, QLP1B1, most of these sequences are in a repetitive region near one end of the erroneous inversion in the female assembly, and their female genetic map positions fall into two sets, consistent with markers within the inversion near its distal end, and ones just distal to the inversion (Supplementary Figure S4). Some of them are at the terminal end of chromosome 12 in the male assembly, which is supported by our female genetic map results; as discussed above, markers in this region of the male assembly generally co-segregate with sex, but their true location could be between 24 and 25 Mb, next to the less terminal region in which the male and female assemblies are syntenic (Supplementary Figure S7). Others are assembled near 11 Mb, and again their female genetic map locations differ, some being consistent with this physical position, while others co-segregate with markers proximal to PAR1 (in the female assembly, some distance away from the boundary), consistent with the possibility that contig IV sequences, or duplicates of them, could include the male-determining factor (Fraser *et al*. 2020).

**Table 1.** SNP positions that are informative about the recombination events in the QHPG5, QLPB1 and PMLPB2 families. The results suggest that the male-determiner is within Region 2a. The positions of the markers are in Mb in the female assembly, where known, because this assembly is annotated, but the positions are mostly very similar in the male assembly, except for the error in the female assembly, where sequences between about 10 and 20 Mb are inverted (Supplementary Table 3E includes positions in both assemblies for the PMLPB2 family, where this is important).

| QHPG5 sibship 2 recombinant male |  |  |  | QLPB1 sibship 2 recombinant male |  |  |  | PMLPB2 sibship 1 recombinant female |  |  |  |
| --- | --- | --- | --- | --- | --- | --- | --- | --- | --- | --- | --- |
| Region (paternal alleles) | First site | Last site | Size of region Mb) | Region (paternal alleles) | First site | Last site | Size of region (Mb) | Region (paternal alleles) | First site | Last site | Size of region (Mb) |
| 1 (X) | 0.13 | 20.60 | 20.47 | 1 (X) | 0.13 | 20.70 | 20.83 | 1 (X) | 1.27 | 11,748,712 <sup>1</sup> | ~ 20 |
| 2a (Y) | 21.05 | 22.90 | 1.85 |  |  |  |  |  |  |  |  |
| 2b (X) | 24.12 | 24.51 | 0.39 | 2 (Y) | 20.93 | 25.38 | 4.45 | 2 (Y) | 22.76 | 25.82 | 3.06 |
| 2c (Y) | 24.85 | 25.38 | 0.53 |  |  |  |  |  |  |  |  |
| PAR1 | 25.52 | 26.40 | 0.88 | PAR1 | 25.52 | 26.43 | 0.91 | No PAR1 markers mapped |  |  |  |

<sup>1</sup> This is close to the end of the error in the female assembly, and the position is near 20 Mb in the male assembly.

The location of a male-determining factor can also be discovered by analysing linkage disequilibrium (LD) of sex-linked molecular variants. Associations with the sex phenotype are consistently detected for SNPs in natural guppy populations, in two chromosome 12 regions proximal to the boundary between PAR1 and the rest of the chromosome, whenever large enough samples from natural populations are analysed (Charlesworth *et al*. 2020a; Fraser *et al*. 2020; Almeida *et al*. 2021; Qiu *et al*. 2022). One region coincides with the region near 21 Mb in the female assembly detected by our recombinants, where the male-determining factor appears to be located, and the other is near 25 Mb.

It is possible that this reflects the presence of two Y-linked loci with balanced polymorphisms in guppies. Under extremely close linkage with the maleness factor, an SA polymorphism can lead to high Y-X differences for molecular variants, creating high diversity within samples of sex-linked sequences, with a lesser signal in the region between the two loci (Kirkpatrick and Guerrero 2014). The guppy Y is not completely non-recombining, but recombination may be infrequent enough to maintain a peak of molecular polymorphism near the male-determining factor, and another near a locus with a sexually antagonistic polymorphism, such as a male coloration factor within PAR2. If so, the completely Y-linked region may be restricted to a small region near 21 Mb in the female assembly.

Coloration factors are not the only Y-X differences expected to cluster near the completely Y-linked region. SA mutations conferring other male-specific benefits should be enriched within a small linkage distance from the male-determining locus, by the sieving effect outlined in the Introduction. Without reliably segregating simple major effect male coloration and other SA factors, it is currently difficult to test this.

The difficulties in studying their genetics also leave it unclear how many Y-linked male coloration polymorphisms are segregating in natural guppy populations. If different Y-linked polymorphisms exist, linkage disequilibria caused by close linkage with the male-determining factor could produce a complicated situation that might make it difficult to pinpoint their locations, as different males would be expected to have different factors, at different distances from the male-determining factor. This could explain the slightly different regions in which molecular markers show associations with the sexes in different populations in the studies just mentioned

Any coloration genes within PAR1 must be physically close to its boundary, because the frequency of recombinants appears to rise immediately distal to the boundary, rather than recombination being infrequent near the boundary (Figure 3B). This is consistent with direct genetic evidence that all partially Y-linked coloration factors appear to be within 10 cM of the M factor (Winge and Ditlevsen 1947). Although a PAR1 location would create selection for closer linkage between the sex-determining locus and the gene with the polymorphic SA alleles, the close linkage that is required for SA polymorphisms to establish (and fixation to be avoided), implies that changes in recombination would probably be undetectably small.

### Sexual dimorphism in expression

Overall, our results suggest that the guppy Y chromosome is not evolving suppressed recombination. This does not imply that SA mutations were unimportant in its evolution. The male-specific expression of coloration traits is an equally interesting consequence of SA selection. Unlike the evolution of recombination rates, polymorphism is not required. As already mentioned, SA mutations may become fixed if their male benefits outweigh disadvantages in females, and any non-rare variants may then evolve male-limited expression, to remove the disadvantages in females (including variants that establish polymorphisms due to diminishing advantages as they become frequent in populations, as mentioned above). The guppy Y does not appear to carry many sequences that are missing from the X, at least not protein-coding genes, based on the current assemblies. Male coloration factors are therefore probably not Y-specific factors, such as Y-linked insertions that would automatically be dominant, as the X would then have no allele (Winge and Ditlevsen 1947). Moreover, at least some coloration factors can be carried on the X after crossing over.

The coloration traits develop at maturity of males, so their expression probably requires testosterone. The coloration mutations’ effects could therefore have been completely or largely male-limited when they first arose, in which case there is no need to consider how male-specific expression evolved. Evolution of male-specificity is also unlikely for mutations that arose in genes with complete or strong sex linkage, because selection against expression in females would be weak if crossovers very rarely produce female carriers. However, complete male-limitation of expression of the initial mutations cannot explain the apparent under-representation of autosomal factors. The common observation of Y-linkage therefore suggests that a selective sieve acted, with autosomal mutations expressed in females being eliminated. Male-specific expression must subsequently have evolved for autosomal factors that are no longer expressed in females, and similar changes may also have affected expression of traits controlled by Y-linked factors. Possibly selection in females acted to prevent expression of coloration traits in the absence of testosterone, and selection in males could also have led to stronger or more reliable expression in the presence of testosterone, perhaps increasing the dominance levels of these polymorphic factors. Such advantageous changes in the two sexes could, respectively, have involved X- and Y-linked cis-acting factors. Trans-acting autosomal factors might also have suppressed expression in occasional females that inherited the mutations, for example ensuring expression of such factors only in the presence of testosterone, which would still allow expression in heterozygous males. In the future, it will be interesting to test whether such changes in sequences, also often involving SA selection, are detectable.

### Data availability

The genotype data and the files from the analysis of repetitive sequences have been deposited in Figshare (https://figshare.com/account/collections/6644282).

## Acknowledgements

This project was supported by European Research Council (ERC) grant number 695225 (GUPPYSEX). We thank colleagues L. Yong, A. Wilson and D.P. Croft at Exeter University for making the guppy families and providing samples from them for genetic mapping, and I.W. Ramnarine and R. Mahabir (University of the West Indies, Trinidad and Tobago) for assistance with the field collection.

## Supplementary Figures

Supplementary Figure S1. Genetic maps of the 22 autosomes in two guppy families from the Aripo river.

Supplementary Figure S2. GC content estimated in 20kb non-overlapping windows of each guppy chromosome. The results were obtained after masking repeats found in the guppy male reference genome assembly, as described in the Methods section.

Supplementary Figure S3. Linkage map of LG16, whose centromere (CEN) is assembled at the right-hand end of the chromosome. **A**. Analysis of intron GC content and change-point suggesting that crossovers occur at a high rate in the 5% closest to the telomere end (TEL). **B**. Linkage analysis using the QLP1B1 family from sire 3, which yielded the largest sibship (See Table S3D), showing the agreement with the GC results, despite SNPs in part of the left-hand end of the assembly not being genotyped.

Supplementary Figure S4. Genetic mapping of the QLP1B1 family. Males and females were analysed separately, and the analysis used all three large sibships. Circles show results for female meiosis, and squares show results for male meiosis. **A** shows the results with the markers positions in the male genome assembly, while **B** shows them using the female assembly positions. Markers in the region that appears to include the male-determining factor (see main text) are indicated in black for female meiosis, and turquoise for female meiosis. It can be seen that these are near close to 21 Mb in the female assembly, near the end of the assembly error (where sequences are inverted in this assembly), and that, in the male assembly they map either to the region where the same sequences are assembled, or to the distal end of chromosome 12.

Supplementary Figure S5.

A. Diagram of Contig IV, based on Supplementary Figure S17 from Fraser et al. (2020) which described the male genome assembly. The positions in the male assembly are shown for each of the microsatellites tested, with arrows indicating their positions in contig IV, or near it. The y axis of the dot plot shows positions in contig XII (near 24 Mb in the male assembly), where the repetitive sequences are also detected. Only the microsatellites in green font amplified in mapping family parents, and all except marker chr12cIV_AC618 gave complex results and could not be mapped, consistent with part of the region being duplicated more distally (see Figure S4).

B. Results of testing markers. The only marker accurately genotyped was chr12cIV_AC618.

Supplementary Figure S6. Genetic maps from dams in the two Aripo river families, showing map locations near the telomere of the X chromosome for several scaffolds that were unplaced in the male or female assemblies.

Supplementary Figure S7. Diagram to explain the possibility that markers assembled in the terminal region of chromosome 12 may belong more proximally, in the PAR2 region that recombines rarely inn male meiosis. A. Map from male meiosis in the family from a Quare river low-predation site, showing only the region distal to 20 Mb in either the male or female guppy genome assembly, for comparison with the female assembly positions (SNPs proximal to 20 Mb almost all segregate as XY alleles in all families studied). B. Diagram of the terminal region of the chromosome, with the two possible interpretations. The terminal markers may be within PAR1 (above) or they may have been mis-assembled, and the set of markers shown in blue in part A may correspond to the gap in these markers just proximal to PAR1 (below).

## Supplementary Tables

Supplementary Table S1. The 11 families used for genetiic mapping. All parents were from Trinidadian guppy natural populations, but the parents of the GLPGrp1 family were from a captive population maintained by Darren P. Croft at the University of Exeter, because no families were raised from wild-caught Guanapo parents. This family had a strongly female-biased sex ratio (see the footnote and the main test). The names of the three families that were included in high-throughput genotyping experiments are showin in bold and in grey cells in column G.

Supplementary Table S2. Summary of information about the guppy chromosomes.

Supplementary Table S3. Genotypes and genetic mapping results for the sex chromosome pair in the different families in the high-throughput mapping experiments, with the mapped markers listed in the order of the sequences in the male assembly. “Mlt” in column B indicates that multiple copies were detected in the male assembly. Sites co-segregating with the male-determining factor (the “XY” pattern) are indicated by blue in the columns with the marker positions, while autosomal segregation is indicated by green, and likely genotyping errors and sites that may be duplicated are indicated by yellow. Progeny genotypes are also individually coloured to indicate conclusions in the four families. These interpretations are based on markers heterozygous in the sire and/or in the dam of the sibship; when both parents have informative genotypes, only homozygotes are used to make conclusions about sex-linkage or autosomal inheritance patterns, including pseudo-autosomal patterns.

Supplementary Table S4. Genetic map location estimates for microsatellite markers distal to 20 Mb in the guppy male and female assemblies, from sires of all families listed in Table S1 (except for the GLPGrp1 family, most of whose progeny were sired by an XX male, see main text). Markers in unplaced scaffolds that map to the sex chromosome are indicated in grey (with locations shown only for those that could be identified in the female assembly); unplaced scaffolds in the female assembly have names starting with NW_, and the numbers of markers genotyped in those that are unplaced in the male assembly are given in the row for the male assembly (details are at the bottom of Table S3A, and the estimated map locations can be seen in Figure S6). For markers proximal to 20 Mb in the assemblies, column B shows the numbers that behaved as completely sex linked in each family. The columns showing results for PAR markers are ordered by their male genetic map positions, with genes in the opposite order from the order in the male assembly (see Figure S3).

Supplementary Table S5. Genes in the region identified as likely to include the male-determining factor (see Supplementary Table S3D), and their positions in the female meiotic map.

Supplementary Table S6. Summary of PAR boundary locations estimated from in male meiosis on the XY pair in all sibships with high-throughput genetic mapping results. For each sibship the location of the most terminal (most distant from the centromere) SNP that co-segregated with the male-determining factor is shown. This estimates the extent of the region that recombines rarely in male meiosis, which is termed the “XY region” or “PAR2” in the main text and Figure 2. The locations of crossovers in male meiosis on the XY pair are also listed.

